## Supplementary Appendix for "Antibody binding reports spatial heterogeneities in cell membrane organization"

**This document contains:**
Supplementary Text
Figures S1-S12
Legends for Movies S1-S3
Supplementary References

**Other supplementary materials for this manuscript include the following:**
Movies S1-S3

### Supplementary Text

### Theory Development

#### Antibody insertion into a semidilute polymer suspension

To describe the crowding-mediated free energy barrier to antibody binding, we invoke the theory of a hard colloid in a nonadsorbing polymer suspension by Louis et al. [1]. We assume that the free energy penalty of inserting the colloidal antibody into the brush is dominated by entropic effects arising from the exclusion of monomer density from the volume occupied by the colloid $V$, as well as the creation of an interface of area $A_\mathrm{s}$ around the colloid. Thus, the free energy can be defined in terms of the osmotic pressure $\Pi$ and interfacial tension $\gamma$.

$$\Delta U = \Pi\left(\phi\right) V + \gamma\left(\phi\right) A_\mathrm{s} \tag{1}$$

##### Brush osmotic pressure

The osmotic pressure can be defined in terms of the monomer volume fraction $\phi$ via the virial equation of state [2]

$$\Pi\left(\phi\right) = \frac{k_\mathrm{B}T}{\nu_\mathrm{p}N}\phi\left(1 + A_2\phi + A_3\phi^2 + \cdots\right) \tag{2}$$

where $\nu_\mathrm{p}$ is the volume of a monomer, $N$ is degree of polymerization, and $A_2$ and $A_3$ are the dimensionless second and third virial coefficients. We operate in the semi-dilute limit with high interpenetration between chains, such that $\phi/\phi^* \gg 1$, where $\phi^*$ is the chain overlap volume fraction. Thus the osmotic pressure can be expressed as

$$\Pi = \frac{k_\mathrm{B}T}{\nu_p N}\phi\left(1 + C_2\frac{\phi}{\phi^*} + \cdots\right) \sim \frac{k_\mathrm{B}T}{\nu_\mathrm{p}N}\phi\left(\frac{\phi}{\phi^*}\right)^m. \tag{3}$$

Upon substituting the scaling $\phi^* \sim N^{-4/5}$, we can express osmotic pressure as

$$\Pi \sim \frac{k_\mathrm{B}T}{\nu_p N}\phi^{m+1}N^{4m/5-1}. \tag{4}$$

Upon assuming that the dominant multi-body interactions between monomers are independent of chain identity, we can impose the constraint that the osmotic pressure must be independent of degree of polymerization. Thus $m = 5/4$ and we achieve the scaling

$$\Pi \sim \phi^{9/4}, \tag{5}$$

which is well-known in the polymer literature [2, 3].

##### Surface tension

In the continuum limit, the antibody diameter is much larger than the monomers such that curvature can be neglected when calculating interfacial tension. Thus we may apply the following relation for interfacial tension near a wall [1]

$$\gamma\left(\phi\right) = -\Pi\left(\phi\right)\Gamma\left(\phi\right) + \int_0^\phi \Pi\left(\phi'\right)\frac{\partial\Gamma\left(\phi'\right)}{\partial\phi'}\mathrm{d}\phi' \tag{6}$$

where $\Gamma$ is the reduced adsorption, defined by the depletion of nonadsorbing hard sphere monomers as a function of distance $r$ from the colloid surface:

$$\Gamma = \int_0^\infty \left(\frac{\phi\left(r\right)}{\phi\left(r \to \infty\right)} - 1\right)\mathrm{d}r. \tag{7}$$

If the monomer density is sufficiently small such that polymer-colloid interactions far outweigh polymer-polymer interactions, then $\Gamma$ will be independent of $\phi$ and the first term of Eq. 6 will dominate. Further assuming an ideal brush gives $\Gamma \approx -2R_g/\sqrt{\pi} \approx R_g$ and reduces Eq. 6 to $\gamma = \Pi(\phi) R_g$. Thus we approximate the interfacial free energy to be the virtual work of creating an cavity of volume $R_g A_s$ around the colloid. Thus we define an effective volume $V^{\text{eff}} = \frac{4}{3}\pi R^3 + 4\pi R^2 R_g$ such that the free energy of including a colloidal antibody into a polymer brush depends solely on the osmotic pressure:

$$\Delta U \approx \Pi(\phi) V^{\text{eff}}. \tag{8}$$

#### Crowding landscape of a polymer brush

We invoke the Milner, Witten, and Cates [4] self-consistent field (SCF) description of the monomer distribution in a semidilute polymer brush, to predict glycocalyx crowding variation with height. In the limit of strong stretching, the monomer volume fraction approximately follows a parabolic form as a function of height $z$ above the grafting surface

$$\phi(z) = \phi_s \left[1 - \left(\frac{z}{L_0}\right)^2\right]. \tag{9}$$

Here $\phi_s$ is the surface volume fraction $\phi(z = 0)$ and $L_0$ is the height at which monomer density reaches approximately zero. For a PEG2k brush, we measured $\langle h \rangle$ and thus set the brush size using the relation $L_0 = 16\langle h \rangle/3\pi$. Substituting the osmotic pressure scaling in Eq. 8 and assuming the exclusion of antibody to scale linearly with osmotic pressure as in Eq. 5, yields a theoretical crowding potential profile

$$\frac{\Delta U}{k_B T} = \Delta U_{0.5k} \left[1 - \left(\frac{z}{L_0}\right)^2\right]^{9/4}. \tag{10}$$

where $\Delta U_{0.5k} = \Delta U(z = 0)$ is the potential at the brush surface, reported by the PEG0.5k sensors in our experiments.

#### Accounting for antigen flexibility

In Fig. S1A, $\Delta U$ represents the repulsive potential experienced by a colloid, like an antibody, inserted into a brush at a single height $z$. However, in our synthetic antigen sensors, the PEG chains linking the FITC antigens to the cholesterol anchors are flexible, enabling FITC to sample a distribution of heights $P_{\text{FITC}}(z)$ with mean height $\langle h \rangle$. In our experiments, for a sensor of a given PEG molecular weight, we measure $\langle h \rangle$ using CSOP and a mean potential $\langle \Delta U \rangle$ that encodes crowding data across the domain of $P_{\text{FITC}}$. Therefore, we weighted the potential distribution predicted by Eq. 10 by $P_{\text{FITC}}$ to predict $\langle \Delta U \rangle$ for PEG sensors of a given $\langle h \rangle$.

$$\langle \Delta U \rangle = \int_0^\infty \Delta U(z) P_{\text{FITC}}(z)\, dz \tag{11}$$

Simulation results showed that the antigen PEG chains behave largely as polymer mushrooms rather than brushes, so we predicted $P_{\text{FITC}}$ using an SCF model for a continuous Gaussian chain [5]. In the case of an ideal, dilute chain with $N$ monomers of length $l$, and with one endpoint tethered at $z = 0$, the density of configurations $G$ for a chain with end-monomer position $z$ is

$$G(z) = \left(\frac{3}{2\pi l^2 N}\right)^{1/2} \left[\exp\left(\frac{-3(z - l/2)^2}{2l^2 N}\right) - \exp\left(\frac{-3(z + l/2)^2}{2l^2 N}\right)\right]. \tag{12}$$

Thus, the normalized FITC probability distribution for a chain of given $l$ and $N$ is

$$P_{\mathrm{FITC}} = \frac{G(z)}{\int_0^\infty G(z)\,\mathrm{d}z} = \frac{\exp\left(\frac{-3(z-l/2)^2}{2l^2N}\right) - \exp\left(\frac{-3(z+l/2)^2}{2l^2N}\right)}{l\sqrt{\frac{2\pi N}{3}}\mathrm{erf}\left(\sqrt{\frac{3}{8N}}\right)}, \tag{13}$$

which is plotted in Fig S1B for PEG ($l = 0.6$nm) [5]. We numerically integrated Eq. 11 across all space for a series of sensors with varying $N$, and therefore $\langle h \rangle$, amongst a PEG2k blocking brush with the potential profile described by Eq. 10. The result is a single weighted potential value that depends on mean sensor height $\langle \Delta U \rangle (\langle h \rangle)$, plotted in Fig. S1C as well as main text Fig. 2B.

##### Red blood cell bidisperse polymer brush

In our theoretical description of red blood cells (RBC), we modified the Milner, Witten, and Cates SCF brush to account for the contributions of two crowding proteins: Glycophorin A (GYPA) and Band 3.

Glycophorin A has 72 disordered extracellular residues with 15 four-sugar O-glycans and one eight-sugar N-glycan [6]. We assume each sugar to be approximately 1 nm in size, and thus approximate the statistical segment length to be of similar size to the side chains, $l = 4$ nm. With 72 amino acids, we approximate the contour length to be $L = 26$ nm, giving $N = 6.5$ statistical segments. For a self-avoiding chain, the corresponding root-mean-squared end-to-end distance $\langle R^2 \rangle^{1/2} = N^{0.588} l = 12$ nm [2]. We thus approximate GYPA to have a mean height $\langle h \rangle = 12$ nm, so that $L_0 \approx 20$ nm. We confirmed this approximation using cell surface optical profilometry [7] to measure the height of an anti-glycophorin A/B antibody that binds to the N-terminus, finding that it too yielded a mean height of 12 nm. Band 3 has a two-pronged N-linked glycan with the longer fork being 19 sugars long, such that $L \approx 19$ nm [6]. We assume the two forks to be of a similar size and consider each glycan to contain two chains of size $N = 19$ with $l = 1$ nm, with a mean height of $\langle h \rangle = 5.6$ nm.

Based on proteomics data, we approximate a GYPA grafting density of 1300/µm$^2$ and a Band 3 density of 6700/µm$^2$ [6, 8, 9]. We estimate the surface volume fraction $\phi_\mathrm{s}$ prefactor in Eq. 9 to be

$$\phi_\mathrm{s} \approx \left(\frac{l^2}{n}\right)^{2/3} \tag{14}$$

where $n$ is the chain grafting density on the surface [3]. Thus, we approximate $\phi_{\mathrm{s,Band3}} \approx 0.070$ and $\phi_{\mathrm{s,GYPA}} \approx 0.075$. Finally, we superimposed the volume fraction profiles given by Eq. 9 for each polymer, to approximate the total volume fraction distribution above the surface to be

$$\phi = \phi_{\mathrm{band3}} + \phi_{\mathrm{GYPA}}, \tag{15}$$

shown in Fig. S2B. By applying the scaling in Eq. 5, we obtain the bidisperse brush potential $\Delta U$ plotted in Fig. S2C.

We account for the flexibility of PEG-FITC sensors by weighting the bidisperse $\Delta U$ by the FITC distribution in Eq. 13 for a series of sensors with mean height $\langle h \rangle$, and then integrating Eq. 11. The weighted brush potential is plotted as a function of FITC mean height in Fig. S2D.

#### The Hill isotherm provides a superior fit for binding data

To determine the dissociation constant $K_\mathrm{D}$, fluorescence intensities $I$ of bound antibody on the bead surface were normalized to a saturation intensity $I_\infty$, and the resulting fraction of bound species $\theta = I/I_\infty$ fit to a binding isotherm as a function of bulk concentration $c_{\mathrm{bulk}}$.

The commonly-used Langmuir isotherm, which assumes monovalent binding and no interaction between bound species, is of the form

$$\theta = \frac{c_{\text{bulk}}}{K_{\text{D}} + c_{\text{bulk}}}. \tag{16}$$

Fig. S3A offers an example of our IgG binding data, which follows a form similar to the Langmuir isotherm at large bulk concentrations, but deviates from the approximately linear, concave-down behavior at low $c_{\text{bulk}}$. We consistently observed this deviation across all of our samples, suggesting polyvalency, cooperative interactions between bound species, or other nonideal effects may warrant the use of a different isotherm.

Polyvalent proteins do not always follow the Langmuir model when binding to ligands on a fluid lipid bilayer, as individual binding events may not occur independently of one another [10]. Cremer et al. demonstrated this effect with cholera toxin B (CTB), a pentavalent protein that binds to ganglioside GM1 [10]. Cremer et al. fit CTB binding data to the Hill isotherm (Eq. 17), which includes a cooperativity exponent $n$ and recovers the Langmuir isotherm in the $n = 1$, finding that $n$ increased with ligand density, ultimately saturating at $n = 2$ [10].

$$\theta = \frac{c_{\text{bulk}}^n}{K_{\text{D}}^n + c_{\text{bulk}}^n} \tag{17}$$

Similar experiments on IgG have also demonstrated a cooperative relationship between the first and second binding events of the two Fab arms, when binding to small molecule haptens on supported lipid bilayers [11, 12]. Cremer et al. [11, 13] further proposed a modified form of the Langmuir isotherm

$$\theta = \frac{\alpha c_{\text{bulk}}}{K_{\text{D}} + c_{\text{bulk}}}. \tag{18}$$

Here $\alpha$ varies with the number of available sites on the surface, defined by the difference between the total concentration of haptens on the surface and the concentration of bound haptens: $c_{\text{s}} = c_{\text{s,total}} - c_{\text{s,bound}}$, and follows the form

$$\alpha = \frac{K_{\text{D},2} + c_{\text{s}}}{K_{\text{D},2} + 2c_{\text{s}}}. \tag{19}$$

In this model, the number of available sites decreases with $c_{\text{bulk}}$, causing $\alpha$ to increase with $c_{\text{bulk}}$, implying positive cooperativity of IgG.

In light of the evidence of polyvalent proteins like IgG binding cooperatively on lipid bilayers, and the qualitative superiority of the Hill fit over the Langmuir isotherm to our data (Fig. S3), we fit our binding data to the Hill isotherm in Eq. 17. We fixed the cooperativity exponent $n = 2$ to ensure standardization between samples and to avoid overfitting the data. While we acknowledge that other cooperative effects may have influenced the shape of the binding isotherm, such as cholesterol-IgG complexes inserting into the bilayer, we consistently observed cooperative behavior across all antigen probes, including DOPE-biotin, from which we expect negligible unbinding into the bulk [14]. Furthermore, we use $K_{\text{D}}^2$ as the fitting parameter in the Hill isotherm, conserving the dimensionality of $K_{\text{D}}$ from the thermodynamic definition of equilibrium for a single particle binding to a surface [15]. We further normalize all dissociation constants by the bare surface value: $K_{\text{D}}/K_{\text{D},0}$, reducing the impact of the exact isotherm used.

As shown in Fig. S3B, the precise form of the isotherm does not affect the result in the main text.

#### Cholesterol sensors slightly favor liquid-disordered domains

We imaged GUVs via epifluorescence microscopy, with 2:2:1 DOPC:DPPC:cholesterol, along with DOPE-rhodamine (0.1%), and cholesterol-PEG0.5k-FITC sensors. The DOPE-rhodamine strongly partitioned into the Ld phase, allowing Ld to be differentiated from Lo. In figure S12 the cholesterol-PEG sensors display a slight preference for the Ld phase over the Lo phase. We hypothesize that the PEG chains disrupt the packing order of the cholesterol, thus causing the sensors to favor Ld while unfunctionalized cholesterol would favor Lo [16]. Thus, the crowding measurements with the cholesterol-PEG0.5k-FITC sensors on HeLa and T47D cells are biased toward the liquid disordered, non-raft phase, but still measure considerable binding in the raft phase.

#### Sensors and antibodies approach equilibrium within an hour

To check that our molecular probes reached equilibrium during the incubation time provided, we measured the intensity of both cholesterol-PEG0.5k-FITC sensors and αCD47 antibodies on red blood cells as a function of time, using a flow cytometer. Fig. S4 shows the increase of IgG and cholesterol-PEG0.5k-FITC signal over one hour, for cells incubated in 100 nM cholesterol and 0.25 µg/mL αCD47 IgG. In our experiments, lipid bilayers were incubated in cholesterol sensors for 30 minutes, ensuring that the sensors reached equilibrium (Fig. S4A). Beads and cells were also incubated in IgG for 30 minutes before collecting image intensity data to find $K_\mathrm{D}$. Fig. S4B suggests that IgG takes approximately one hour to fully reach equilibrium in a flow cytometer. In our experiments, however, we qualitatively found that when allowed to sediment under gravity in the imaging chamber, and particularly when subjected to light pipette mixing, beads and cells reached equilibrium much more quickly. We thus infer that IgG binding kinetics are transport-limited, and that due to the advection introduced by mixing and settling under gravity, our beads and red blood cells were approximately equilibrated. Mammalian cells were incubated in IgG for one hour, ensuring equilibration.

#### MD Simulation Details

##### Coarse-grained model

The prepared systems are described using the overdamped Langevin equations of motion, also known as Brownian dynamics, where the velocity $\dot{\mathbf{x}}_i = \mathbf{F}_i/\gamma_i$ of particle $i$ is numerically integrated forward in time and $\mathbf{F}_i$ is the sum of all forces on particle $i$. All simulations were performed using the GPU-enabled HOOMD-blue simulation package [17].

We coarse-grain (CG) PEG molecules as Kremer-Grest bead-spring polymer chains according to [18], where each CG bead represents the C-O-C monomer unit with length $\sigma = 0.33$ nm. Although our coarse-graining is at an atomic scale, we will still assume an implicit solvent in this simplified model and neglect any polymer interactions with the surrounding solvent molecules.

Non-bonded interactions between monomer pairs are modeled via a purely repulsive Weeks-Chandler-Anderson (WCA) potential

$$V_{\mathrm{WCA}}(\mathbf{r}) = \begin{cases} 0 & \text{if } |r| \geq 2^{1/6}\sigma \\ 4\epsilon\left(\left(\frac{\sigma}{|r|}\right)^{12} - \left(\frac{\sigma}{|r|}\right)^{6}\right) & \text{if } |r| < 2^{1/6}\sigma \end{cases} \tag{20}$$

where we set $\epsilon = k_\mathrm{B}T$.

Bonded monomers along the polymer chain interact through the "finite extensible nonlinear elastic" (FENE) potential:

$$V_{\mathrm{FENE}}(\mathbf{r}) = \frac{1}{2}kr_0^2 \ln\left(1 - \left(\frac{\mathbf{r}}{r_0}\right)^2\right) + V_{\mathrm{WCA}}(\mathbf{r}). \tag{21}$$

Additionally, the flexibility of polymer chains is represented using a harmonic bending potential:

$$V_{\mathrm{harm}}(\theta) = \frac{l_{\mathrm{p}}}{2l_{\mathrm{b}}}(\theta - \theta_0)^2 \tag{22}$$

where $\theta$ is the angle between three monomers along a chain. We choose the persistence length $l_{\mathrm{p}} = \sigma$ and equilibrium bond length as $l_{\mathrm{b}} = \sigma$.

All surface polymers are tethered using two wall-potentials $V_{\mathrm{wall}}$ which constrains the vertical position of the first monomer, similar to MD methods in [7].

Antibodies interact through a similar WCA potential with PEG monomers:

$$V_{\mathrm{WCA}}(\boldsymbol{r}) = \begin{cases} 0 & \text{if } |r| \geq 2^{1/6} r_{\mathrm{a}} \\ 4\epsilon\left(\left(\frac{r_{\mathrm{a}}}{|\mathbf{r}|}\right)^{12} - \left(\frac{r_{\mathrm{a}}}{|\mathbf{r}|}\right)^{6}\right) & \text{if } |r| < 2^{1/6} r_{\mathrm{a}} \end{cases} \tag{23}$$

where the equilibrium distance is $r_{\mathrm{a}} = (\sigma + d_{\mathrm{a}})/2$ where $d_{\mathrm{a}}$ is the coarse-grained antibody diameter.

#### Average sensor height

We simulated sensor polymers at a surface density of $1000/\mu\mathrm{m}^2$, in agreement with the experimental coverage of PEG-FITC conjugates on silica beads.

The degree of polymerization was varied from $N = 10$ to $N = 200$ to span the range of polymer contour lengths between PEG0.5k and PEG10k. After an initial equilibration period, we track the vertical height $z$ of the end monomer above the surface, which corresponds to the location of the FITC sensor. The monomer hard sphere radius $r_{\mathrm{HS}} = 2^{1/6}(\sigma/2)$ was subtracted from $z$ since the hard sphere bead never overlaps with the underlying surface. The spatial data were binned and averaged to compute the probability distribution $P(z)$ of sensor positions for various polymer linker lengths. The relationship between $N$ and $\langle h \rangle$ is observed to follow $\langle h \rangle \sim N^{3/5}$, in agreement with Flory theory of a mushroom brush (Fig. S5). This result is expected given the diluteness of the polymer sensors on the surface.

#### Antibody insertion into PEG brush

We simulate free antibody with PEG2k polymers with $N = 45$ tethered to the cell surface at a surface density of $30000/\mu\mathrm{m}^2$. After equilibration, the center of mass position of the antibody was binned as a function of height above the surface, and the hard sphere radius of the antibody was again subtracted such that $z = 0$ indicates antibodies that are flush with the cell surface. The size of an IgG antibody is $\approx 10$ nm [19, 20], which is larger than the PEG2k brush size of $\approx 2 - 3$ nm. We assume that only the Fab region sticks into the PEG2k brush. Therefore, we modeled only the Fab region of the IgG with 4 nm spherical particles, which also recovered he experimentally observed brush potential of $\Delta U = 1k_{\mathrm{B}}T$ at the surface. To obtain the effective brush crowding potential $\langle \Delta U \rangle$, numerical integration of Eq. 11 was performed and normalized by the effective potential of the PEG0.5k sensor.

#### Coarse graining RBC proteins

Based on proteomics literature, we model the RBC cell surface as a bidisperse polymer brush consisting of the two most abundant proteins based on extracellular size and surface density, GYPA and Band 3 (Fig. S6). We choose the coarse-grained bead diameter of GYPA to be 4 nm, representing the 4-sugar side chain, and use $N = 7$ beads to maintain the contour length of GYPA to be roughly 28 nm. For Band 3, we coarse-grain the branched N-glycan on the extracellular domain into a single chain with 2 nm beads while choosing $N = 10$ to maintain the $\approx 20$ nm contour length. In doing so, we have neglected the entropic penalty of confining these side branches, which may lead to slight underestimation of the crowding penalty. Since the RBC surface proteins are taller than the size of an IgG, we assume that the whole IgG interacts with the glycocalyx. Therefore, we modeled the IgG with 11 nm spherical particles, which recovered a repulsive penalty of $\Delta U = 2k_\mathrm{B}T$ at the surface, which is reasonable since the RBC brush is taller than the PEG brush and the bulk size of the antibody will dictate the repulsive potential. The effective brush potential $\langle \Delta U \rangle$ was then obtained similarly as in the PEG brush simulations.

Although we have coarse-grained the IgG into a spherical bead, in reality it has a Y-shaped structure consisting of two Fab and one Fc regions. Thus, one potential drawback to simulating the IgG as a spherical particle is the overestimation of the free energy penalty when inserted into the brush. Furthermore, only the Fab regions target and bind to the antigen, so our measured IgG affinity could deviate from experiments. However, our main simulation result is the brush crowding potential ΔU, and we do not expect the IgG coarse-graining to have significant consequences other than a small overestimation of the osmotic penalties associated with insertion.

#### Materials

1,2-dioleoyl-sn-glycero-3-phosphocholine (DOPC, 850375C), 1,2-dioleoyl-sn-glycero-3-phosphoethanolamine-N-[methoxy(polyethylene glycol)-2000] (18:1 PEG2k PE, 880130C), 1,2-dioleoyl-sn-glycero-3- phosphoethanolamine-N-(cap biotinyl) (18:1 biotinyl cap PE, 870273C), and 1,2-dipalmitoyl-sn-glycero-3-phosphocholine (DPPC, 850355C) in chloroform and 1,2-dipalmitoyl-sn-glycero-3-phosphoethanolamine-N-(cap biotinyl) (16:0 biotinyl cap PE, 870277P) in solid form, were purchased from Avanti Polar Lipids (Alabaster, AL).

Cholesterol (stabilized with α-Tocopherol, catalog number: C3624, lot number: YB46F-QS) was purchased from TCI chemical.

Monoclonal antibody raised against Fluorescein (1F8-1E4; catalog number: 31242) was purchased from Invitrogen. Alexa Fluor 647 labeled anti-biotin monoclonal antibody (BK-1/39; catalog number: sc-53179 AF647) was purchased from Santa Cruz Biotechnology. Human HER2/ErB2 C-terminal poly-His protein (catalog number: 10004-H08H), Human CD45 C-terminal poly-His protein (extracellular domain; catalog number: 14197-H08H), and anti-HER2 monoclonal antibody (catalog number: 10004-MM03) were purchased from Sino Biological US. Recombinant NS0-derived mouse CD45 protein with C-terminal 6-His tag was purchased from R&D systems (catalog number: 114-CD-050 Lot NHF0318121). Monoclonal antibodies against mouse CD45RB (C363-16A; catalog number: 103311) and pan-CD45 (I3/2.3; catalog number: 147715) were purchased from BioLegend. Purified anti-human Glycophorin AB monoclonal antibody (CD235ab) (HIR2; catalog number: 306602) was also purchase from BioLegend.

Annexin V, FITC (catalog number: 29001) was purchased from Biotium.

Silica microspheres (4.07µm; catalog code: SS05002; lot number: 12602) were purchased from Bangs Laboratories.

Cholesterol-PEG-Amine, MW 1k (catalog number: PLS-9961), MW 2k (catalog number: PLS-9962), MW 5k (catalog number: PLS-9964), and MW 10k (catalog number: PLS 9965) were purchased from Creative PEGWorks. NHS-Fluorescein (5/6-carboxyfluorescein succinimidyl ester) (NHS-FITC; catalog number:46409) and Zeba Spin Desalting Columns, 7K MWCO (catalog number: 89882) were purchased from Thermo Scientific.

Alexa Fluor 647 NHS Ester (Succinimidyl Ester) (NHS-AF647; catalog number: A20006), Alexa Fluor 555 NHS Ester (Succinimidyl Ester) (NHS-AF555; catalog number: A20009), and Alexa Fluor 488 NHS Ester (Succinimidyl Ester) (NHS-AF488; catalog number: A20000) were purchased from Invitrogen.

Neuraminidase from *Clostridium perfringens* (catalog number: N2876-25UN) was purchased from Sigma.

Trypsin-EDTA (0.05%; catalog number: 25300054), High-glucose Dulbecco's Modified Eagle Media (DMEM; catalog number: 10566024), high-glucose Roswell Park Memorial Institute 1640 media (RPMI; catalog number: 61870036), Penicillin-streptomycin (catalog number: 15140122), and fetal bovine serum (FBS; 10437028) were purchased from Gibco.

Single-donor human whole blood (catalog number: IWB1K2E10ML) was purchsed from Innovative Research. Blood was de-identified. The researchers involved in this study did not take part in sample collection, and did not have any contact with the donor.

MATLAB educational license was obtained from MathWorks Inc.

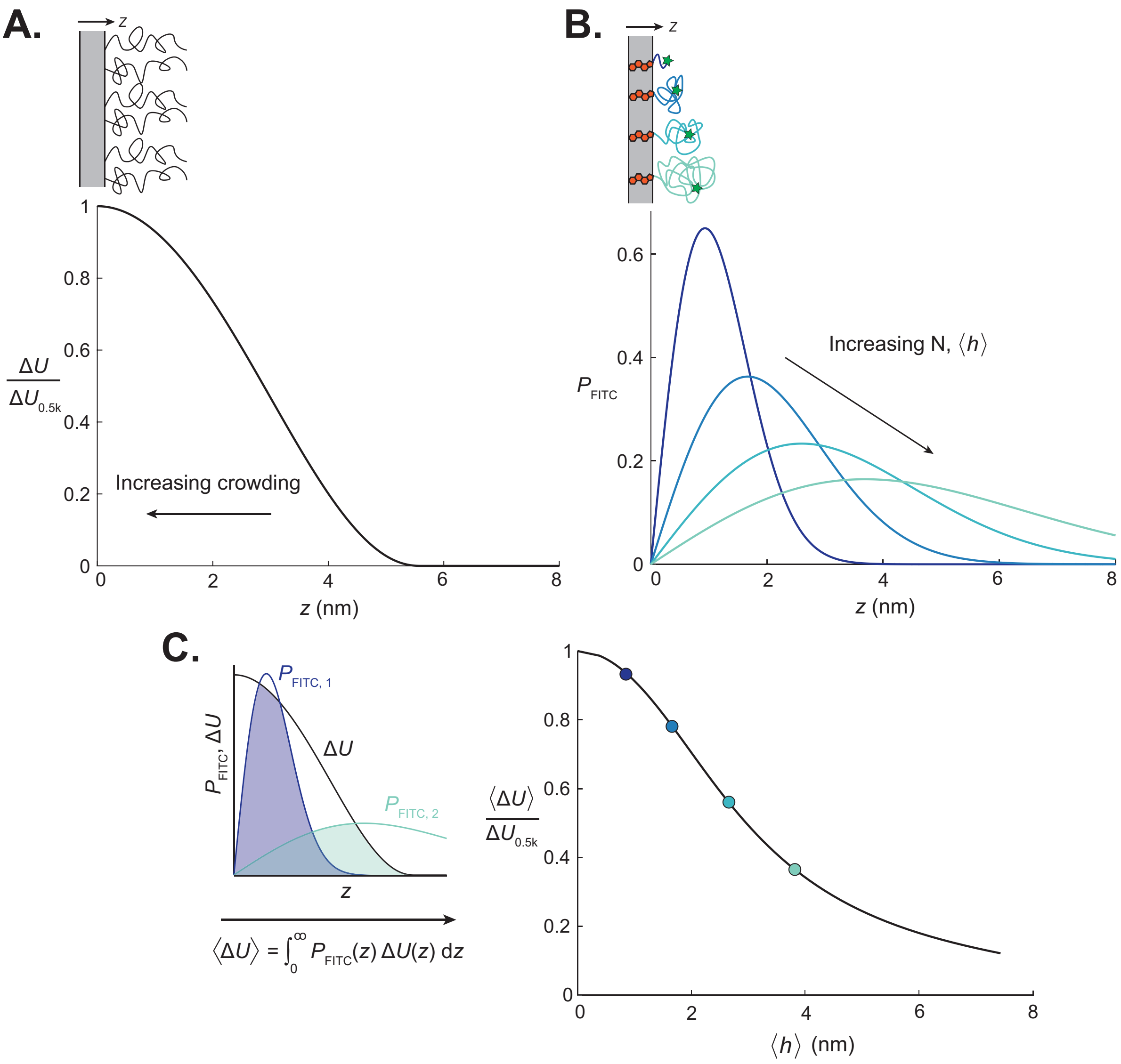


Figure S1: Modeling the crowding landscape experienced by antibodies binding to PEG-tethered FITC antigens in a PEG polymer brush. (A) Repulsive potential $\Delta U$ exerted on an adsorbing colloid by a PEG brush with mean height $\langle h \rangle = 3.3$nm. The repulsive potential varies with distance from the surface $z$ according to the Milner, Witten, and Cates description of monomer density a polymer brush, with the semidilute scaling relationship between osmotic pressure and volume fraction (Eq. 10). The repulsive potential is normalized by the value at the grafting surface, as reported by a PEG0.5k-FITC sensor, $\Delta U_{0.5k}$. (B) End-monomer distribution for surface-tethered polymer mushrooms. Cholesterol-PEG-FITC sensors are modeled as continuous Gaussian chains, with the end monomer, bound to FITC, sampling a distribution of heights $P_{\mathrm{FITC}}$ according to Eq. 13. Curves are plotted for sample polymers with $N = 7, 23, 56$, and 113, with PEG $l = 0.6$ nm. (C) Weighted repulsive potentials account for antigen flexibility and measurement limitations. (Left) Repulsive potentials are weighted by the FITC distribution for each theoretical PEG sensor and integrated across all space to yield a mean potential $\langle \Delta U \rangle$. (Right) Mean potentials are plotted for a series of theoretical sensors, each having a distinct mean height $\langle h \rangle$. Points corresponding to the mean heights of the sample sensors in Fig. S1B are highlighted.

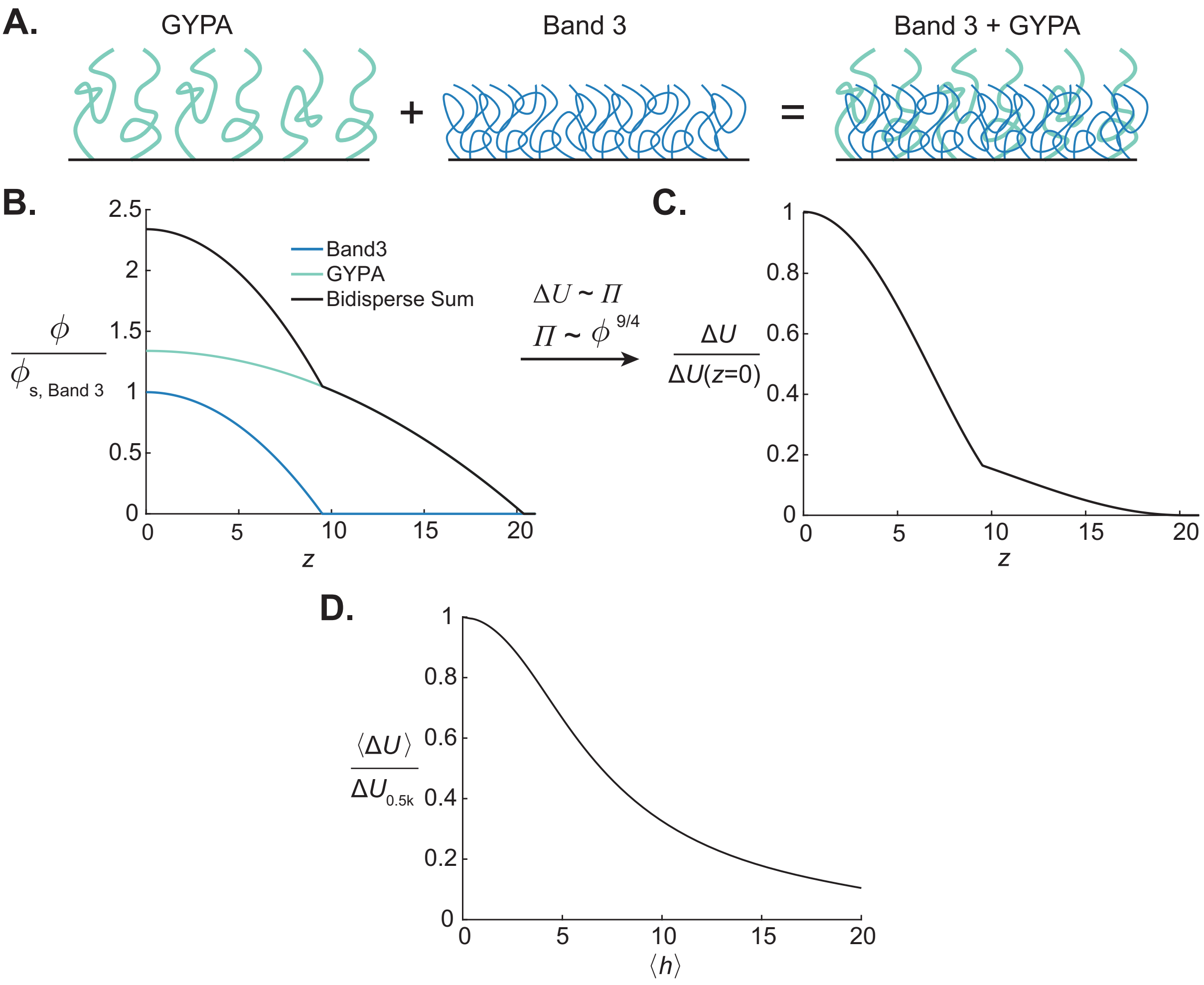


Figure S2: Modeling the red blood cell (RBC) glycocalyx as a bidisperse polymer brush containing glycophorin A (GYPA) and Band 3, using analytical theory. (A) Illustration of the bidisperse polymer brush represented in this analysis. GYPA is taller than Band 3 and has larger monomers but is expressed at a lower density on the RBC surface. Brushes are directly superimposed and assumed to behave independently of one another. (B) Monomer volume fraction distributions according to Eq. 9 are plotted for both GYPA and Band 3 and normalized by the surface volume fraction of Band 3. For each protein, the inputs $\phi_s$ and $L_0$ are estimated based on proteomics data and glycosylation characterization. The profiles for the two proteins are superimposed to produce a bidisperse volume fraction profile. (C) Crowding potential of the glycocalyxis plotted as a function of height. The monomer density profile in Fig. S2B is converted to a repulsive potential that excludes adsorbing IgG using Eqs. 8 and 5. The potential is normalized against its value at the membrane surface. (D) Mean brush potential experienced by PEG sensors on the RBC surface. The bidisperse polymer brush potential $\Delta U$ is weighted by the FITC distribution $P_{\mathrm{FITC}}$ from Fig. S1B for a series of sensors, according to Eq. 11. The resulting mean potential $\langle \Delta U \rangle$ is plotted as a function of mean sensor height $\langle h \rangle$

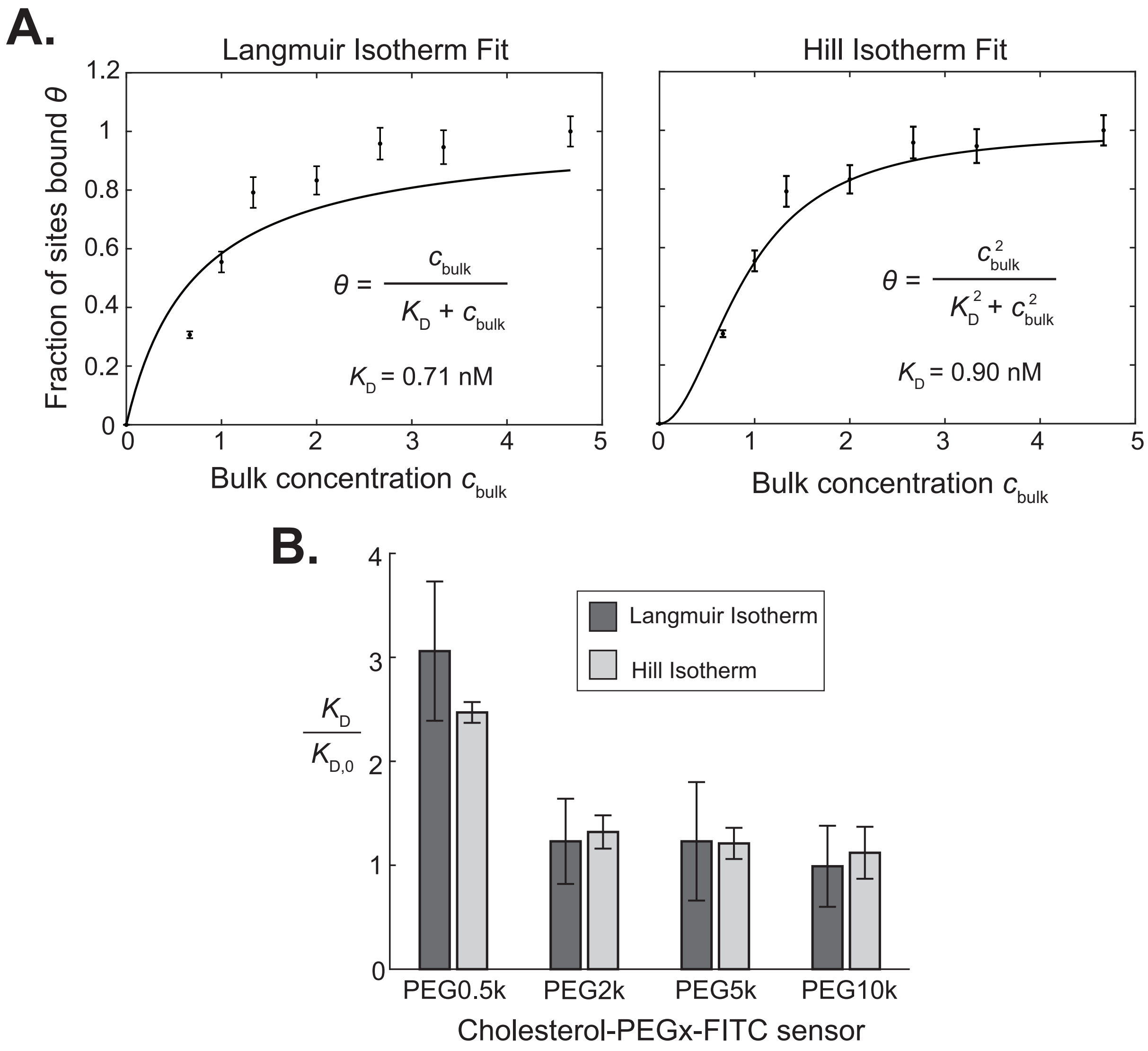


Figure S3: A comparison of Langmuir isotherm and Hill isotherm fits to antibody binding data shows that the fitting method has little impact on our conclusions. (A) For a cholesterol-PEG0.5k-FITC sensor, the fraction of bound antigen sites, normalized by the concentration of antibodies bound at saturation, is plotted as a function of bulk antibody concentration. (Left) The Langmuir isotherm is fit to the data. (Right) The Hill isotherm with cooperativity coefficient $n = 2$ is fit to the data. (B) For all cholesterol-PEGx-FITC sensors on beads we plot the ratio of $K_{\mathrm{D}}$ with a PEG2k brush blocking binding to $K_{\mathrm{D},0}$ with no brush. Dissociation constants were fitted to binding data with both the Langmuir isotherm and Hill isotherm. For all sensors, the ratio of dissociation constants varies by approximately 15% or less, with no significant qualitative difference in the overall trend.

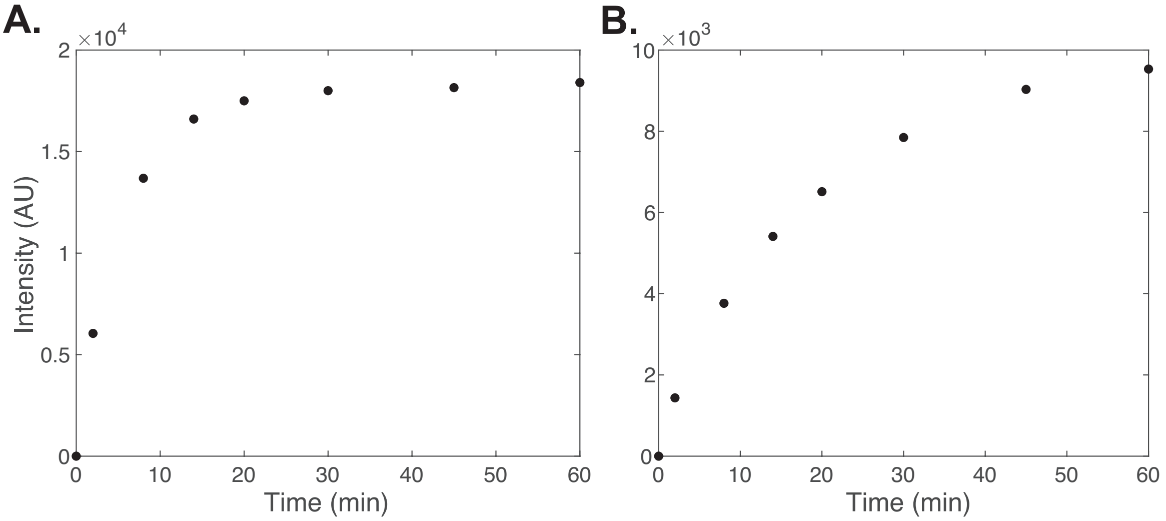


Figure S4: Kinetics of sensor and IgG binding to red blood cells. (A) Red blood cells were incubated in 100 nM cholesterol-PEG0.5k-FITC sensors and the FITC fluorescence intensities measured using flow cytometry. Intensities are plotted as a function of time. (B) Red blood cells were incubated in 0.25 µg/mL αCD47 IgG antibody labeled with Alexa Fluor 647 and the antibody fluorescence intensities plotted as a function of time.

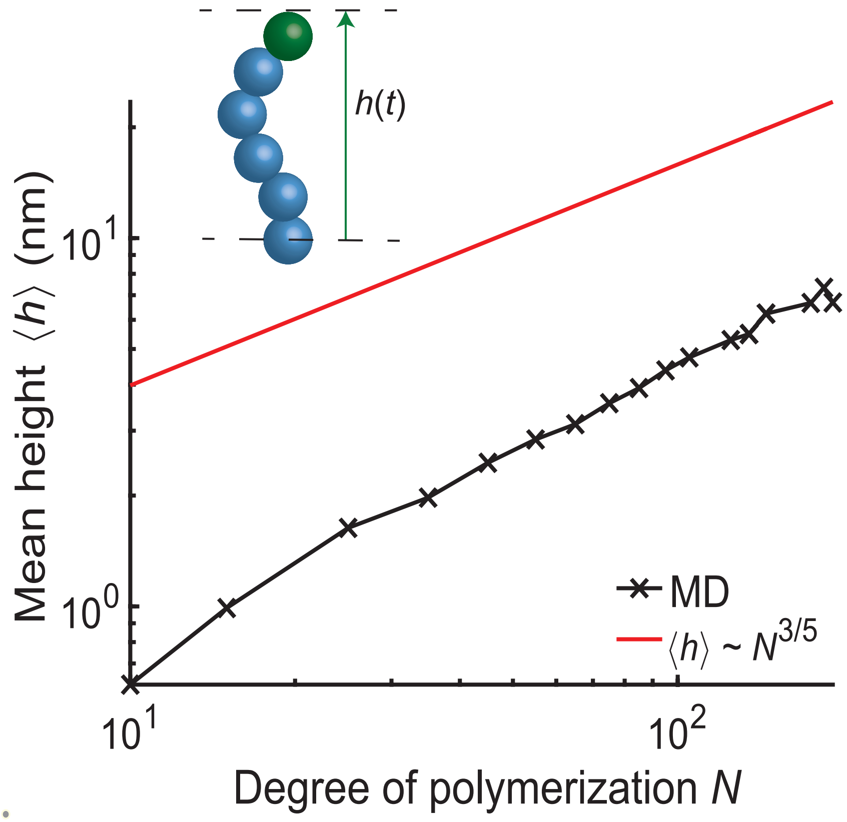


Figure S5: MD simulation of average height of FITC sensor. The average height of the FITC sensor (green) was obtained for various polymer degrees of polymerization, which is controlled by tuning the number of linker monomers (blue). The red theory line corresponds to a self-avoiding swollen polymer chain in a good solvent.

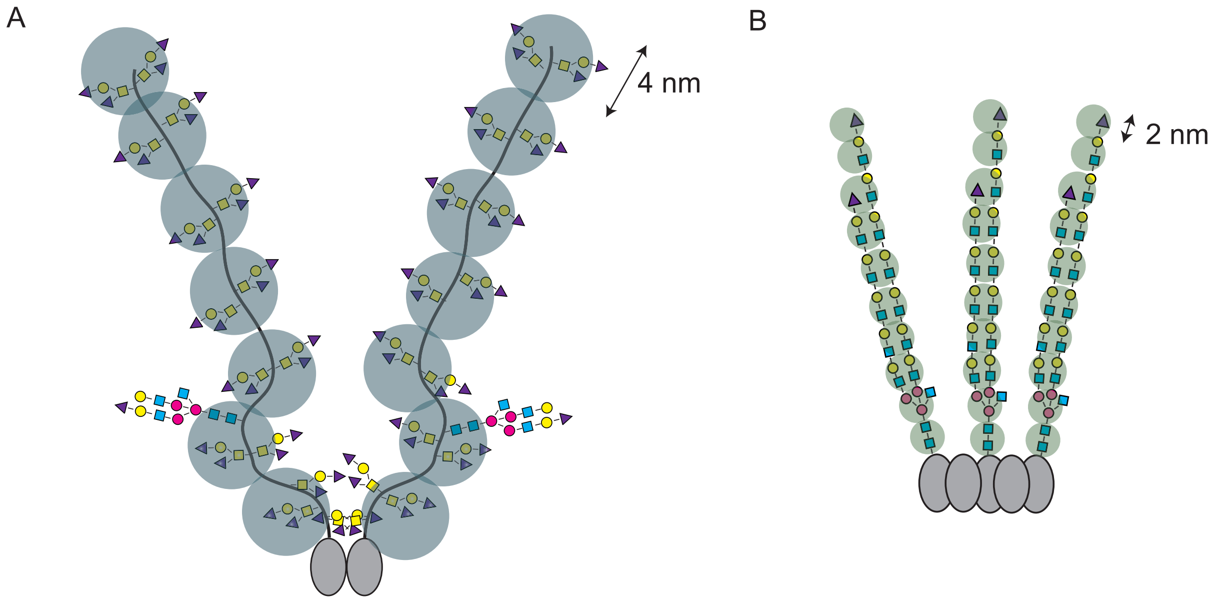


Figure S6: Coarse-graining of Glycophorin A (GYPA) and Band 3 on RBC glycocalyx. (A) In the molecular dynamics (MD) simulations, the extracellular domain of GYPA is modeled as two seven-bead polymer chains with bead diameter of 4 nm corresponding to the length of the sugar side chains. (B) Band 3 is modeled as three ten-bead polymer chains with bead diameter of 2 nm corresponding to a pair of sugars across the two side branches [6].

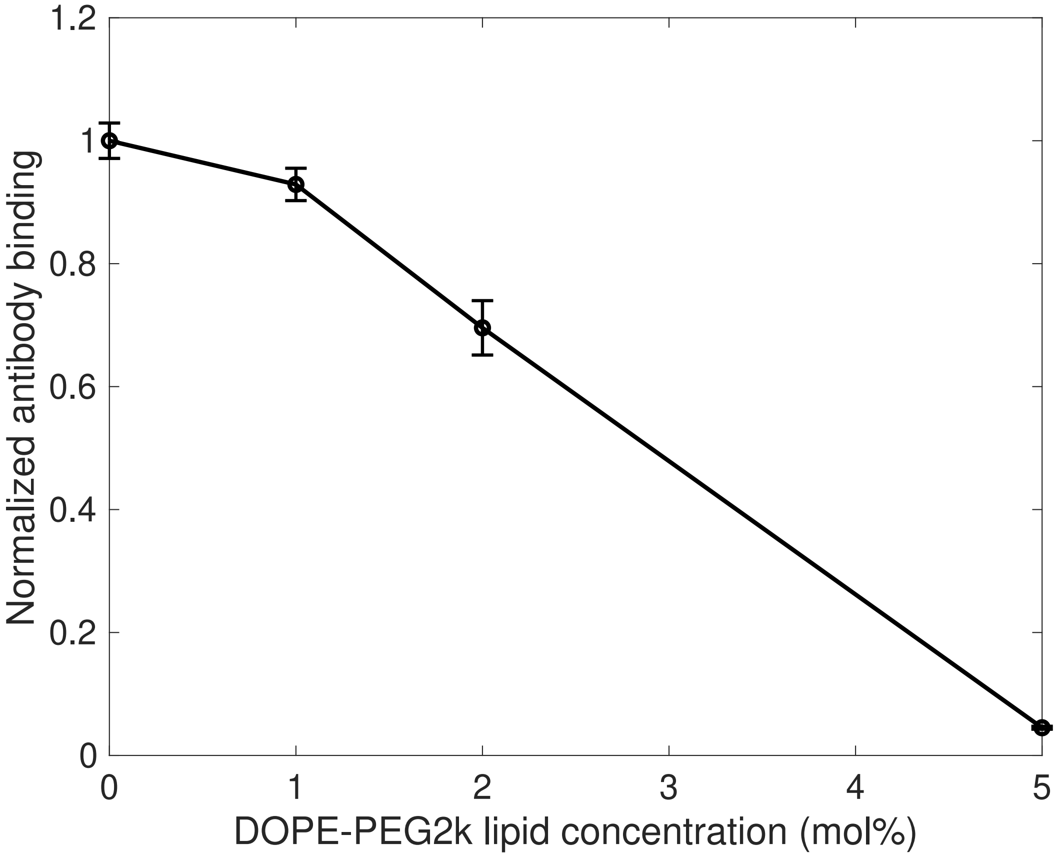


Figure S7: Determining the appropriate PEG brush density for crowding measurements on reconstituted lipid bilayers. Lipid bilayer-coated silica beads containing 0.05% DOPE-FITC and varying lipid fractions of DOPE-PEG2k were incubated in anti-FITC IgG (1 µg/mL) for 30 minutes. The fluorescence intensity of Alexa Fluor 647-labeled anti-FITC is plotted here, normalized to the bare value. Interpolation shows an expected antibody binding drop of approximately 50% compared to the bare surface for 3% DOPE-PEG2k. To ensure sufficient antibody binding for reliable measurements, while still achieving an appropriate decrease for crowding measurements, we chose to use 3% PEG2k in our main text experiments with reconstituted bilayers on beads.

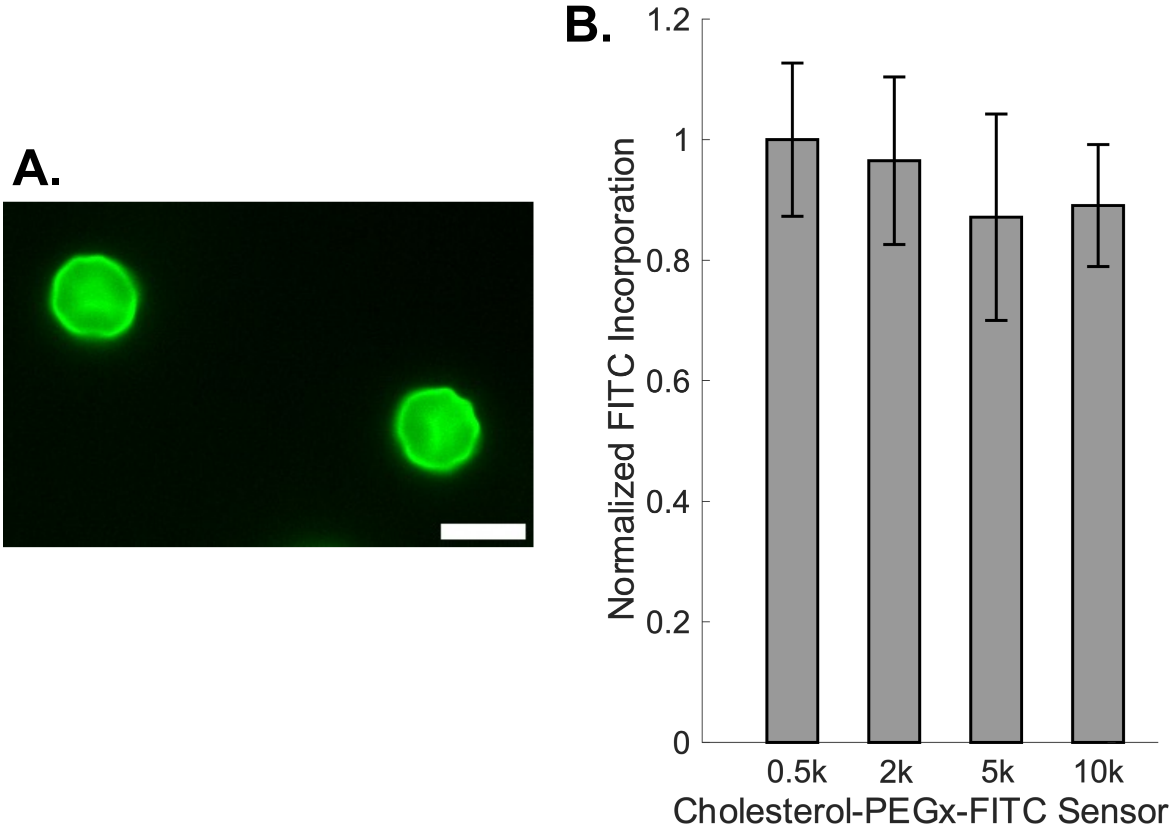


Figure S8: Cholesterol-PEGx-FITC sensor insertion into red blood cell membrane. (A) Representative fluorescence image of FITC on the red blood cell surface. Scale bar is 10 µm. (B) Intensities of FITC on red blood cells incubated with cholesterol-PEGx-FITC (x=0.5k, 2k. 5k, and 10k)are plotted, normalized to that of cholesterol-PEG0.5k-FITC. Antigen densities were equal across experiments to within 15%. Error bars represent standard deviation of n=10 red blood cells.

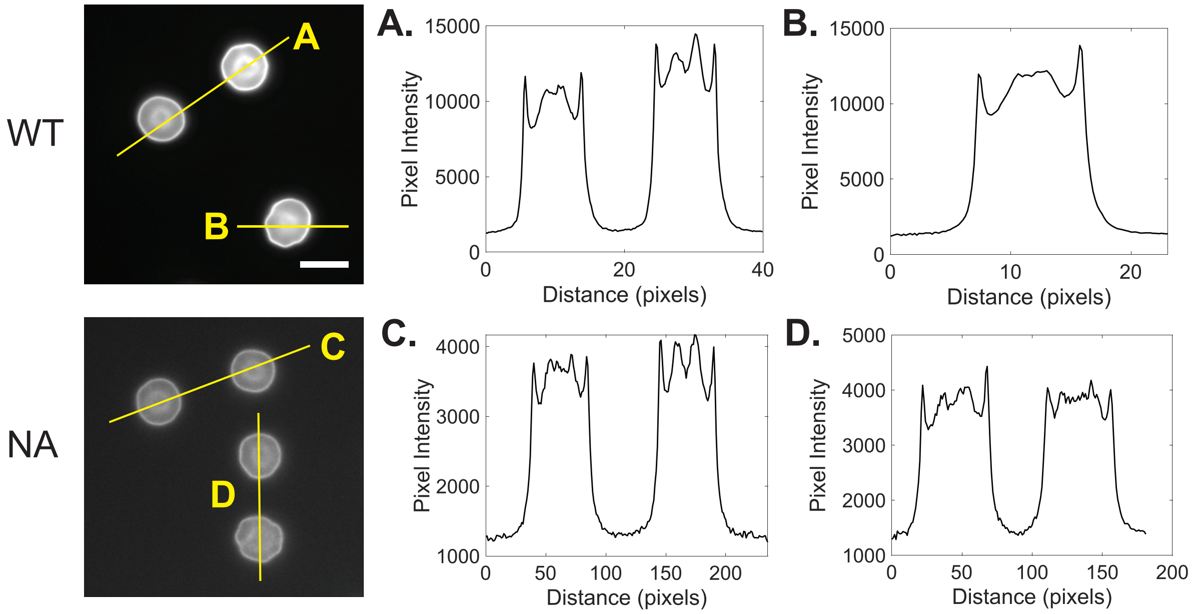


Figure S9: Wheat germ agglutinin (WGA) binding confirms sialic acid removal from red blood cell (RBC) surface. RBCs were treated with 100 mU/mL neuraminidase (NA) for two hours at 37°C to remove surface sialic acid from the glycocalyx. Cells were treated with 0.1 µg/mL wheat germ agglutinin (WGA) for 30 minutes, then imaged to measure binding. WGA is a lectin that binds to sialic acid and N-acetyl-D-glucosamine (GlcNac, a minor constituent of RBC extracellular glycans). (*Left*) Here we show representative images of wild type (WT) RBCs (top) and RBCs treated with NA (bottom), labeled with WGA-Alexa Fluor 647. (*Right*) Line scans (A) and (B) show representative WGA fluorescence intensities on WT RBCs while line scans (C) and (D) show representative WGA fluorescence intensities on NA-treated RBCs. A $\sim 2/3$ reduction in WGA binding upon treatment with NA is apparent, and is indicative of a significant reduction in glycocalyx sialic acid content.

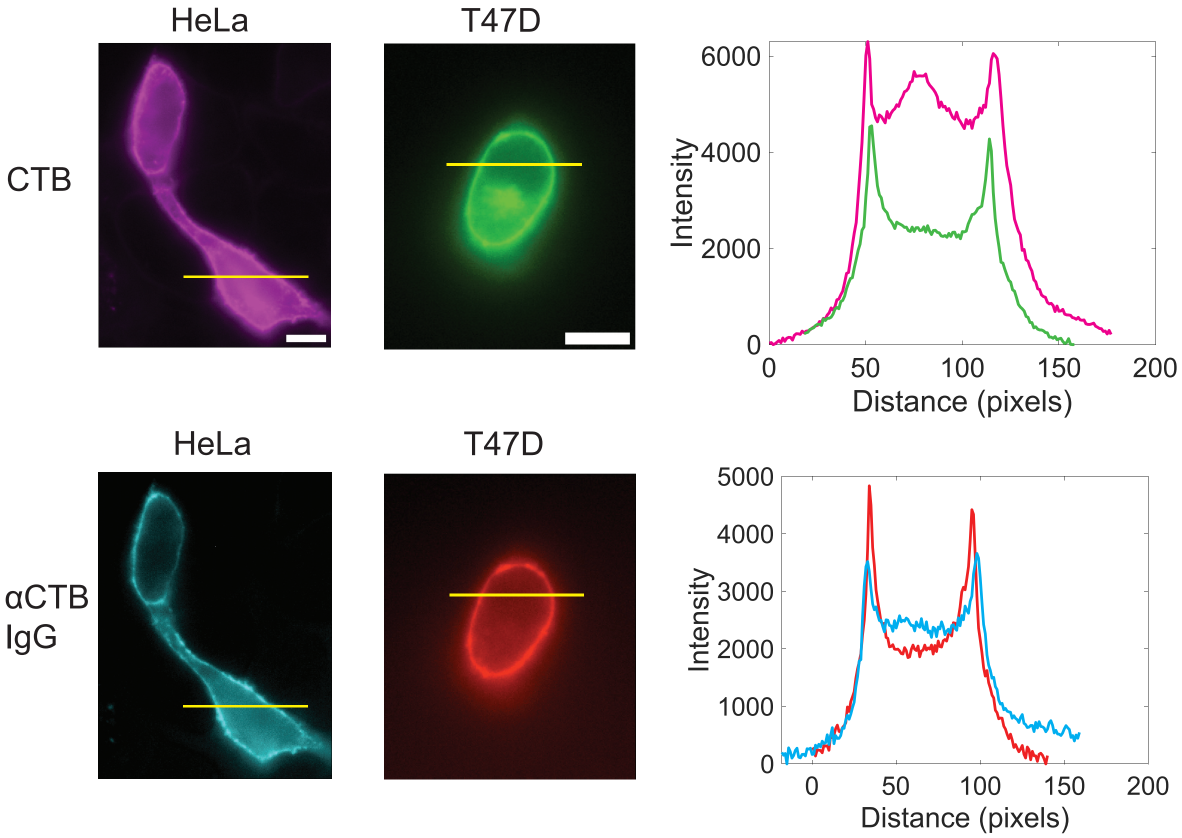


Figure S10: Representative line scans showing cholera toxin B (CTB) and anti-CTB IgG monoclonal antibody (αCTB IgG) fluorescence intensities on HeLa (left) and T47D (right) cells. Yellow lines indicate the position of line scans. Scale bars are 10 μm.

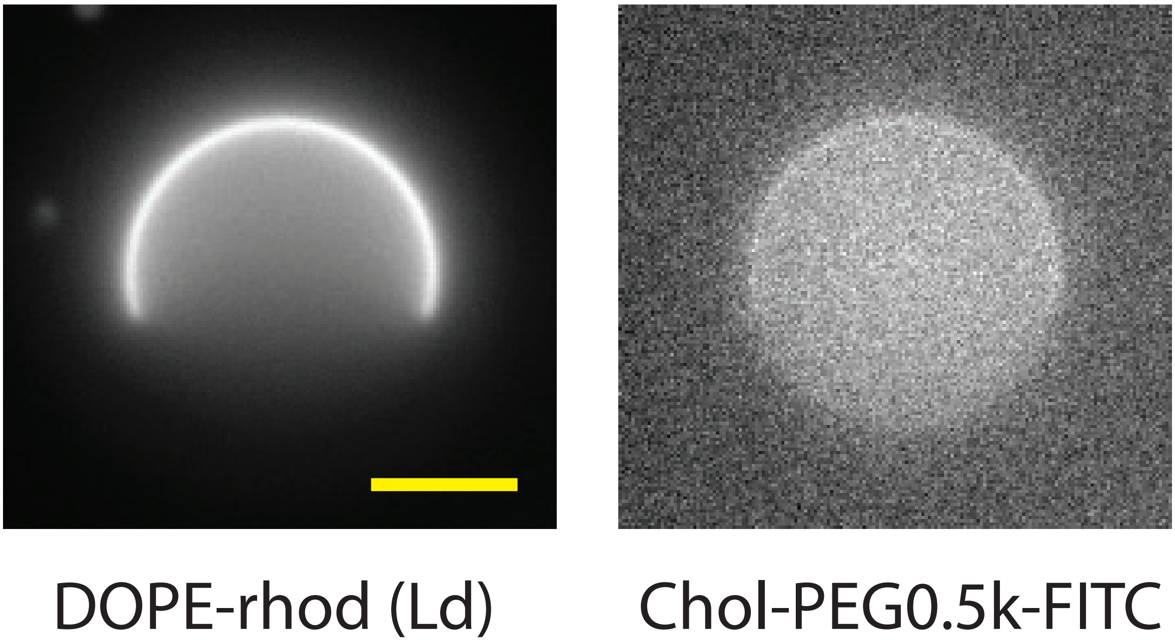


Figure S11: The 555 nm DOPE-rhodamine channel is shown on the left, with the Ld phase of a GUV appearing as a bright crescent. The 488 nm cholesterol-PEG-FITC channel is shown on the right, with a slightly brighter half aligning with the rhodamine channel. Scale bar is 5 µm.

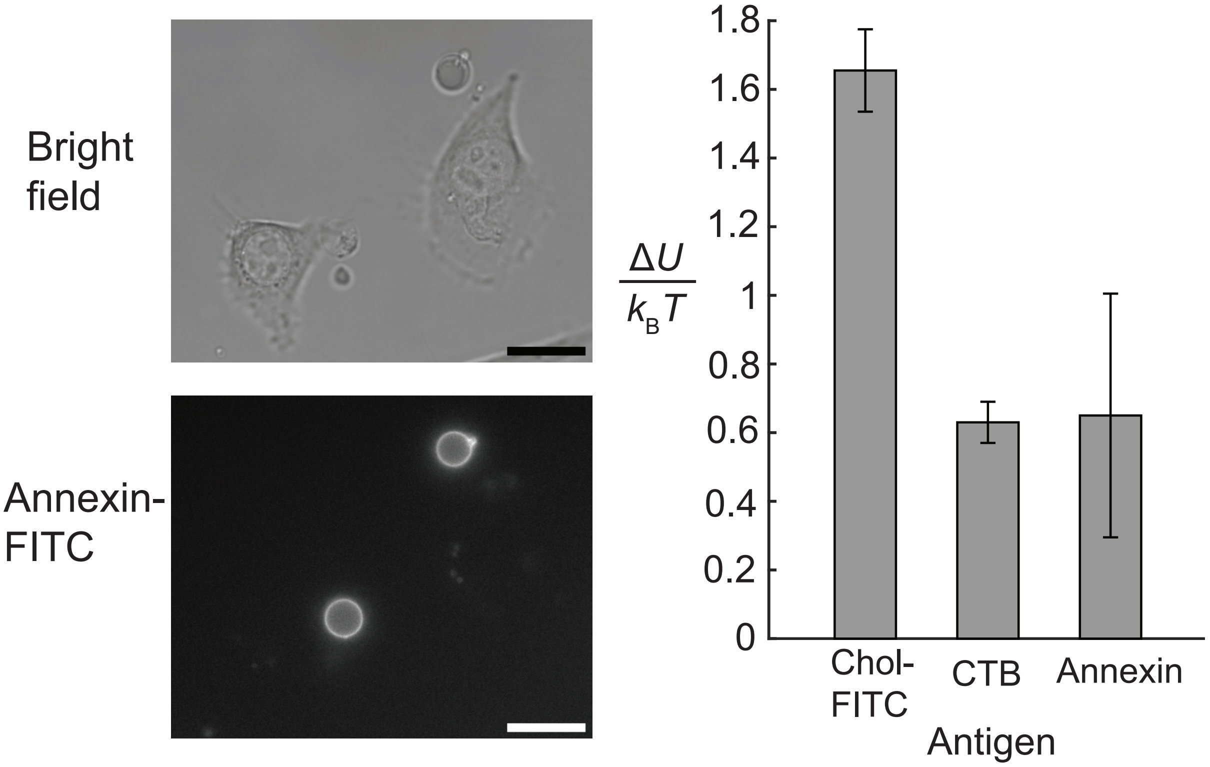


Figure S12: Control to evaluate the role of protein clustering in inducing cell surface heterogeneity. Human cervical cancer HeLa cells were incubated in annexin V-FITC, before being washed and the $K_D$ of αFITC IgG measured. Annexin V binds to phosphatidylserine (PS), which is not typically expressed on the surface of healthy cells. (*Left*) Bright field images of cells are contrasted with fluorescence images of annexin V-FITC. Annexin V-FITC preferentially binds to membrane blebs and vesicles, rather than the cell membrane. Scale bars are 10 µm. (*Right*) The free energy penalty to antibody binding, imposed by the glycocalyx, is plotted for cholesterol-PEG0.5k-FITC (abbreviated chol-FITC), cholera toxin B (CTB), and annexin V antigens. Cholera toxin and annexin V show reduced local crowding relative to cholesterol-PEG0.5k-FITC, possibly as a consequence of their clustering behavior excluding other bulky proteins.

#### Movie Legends

##### Movie S1. MD simulation of FITC sensors

MD simulation of FITC sensor (green) with 64 linker monomer units (blue), each of diameter 0.33 nm. Sensors were tethered to the bottom of the simulation box at a surface density of $1000/\mu m^2$.

##### Movie S2. MD simulations of antibodies in monodisperse PEG surface

MD simulation of ideal gas antibodies (red) of diameter 4 nm and $N = 45$ PEG brush (grey) with monomer diameter 0.33 nm and surface density of $30000/\mu m^2$.

##### Movie S3. MD simulation of antibodies in bidisperse red blood cell surface

MD simulation of ideal gas antibodies (red) of diameter 11 nm in the presence of a bidisperse brush. Glycophorin A (dark green) is represented as a 7 bead chain with monomer diameter 2 nm and surface density of $1300/\mu m^2$, while Band 3 (light green) is represented as a 10 bead chain with monomer diameter 1 nm and surface density of $6700/\mu m^2$.

#### References


(1) Louis, A. A.; Bolhuis, P. G.; Meijer, E. J.; Hansen, J. P. *The Journal of Chemical Physics* **2002**, *116*, 10547–10556.

(2) Rubinstein, M.; Colby, R., *Polymer Physics*, 1st ed.; Oxford University Press: 2003.

(3) Halperin, A. *Langmuir* **1999**, *15*, 2525–2533.

(4) Milner, S. T.; Witten, T. A.; Cates, M. E. *Macromolecules* **1988**, *21*, 2610–2619.

(5) Russel, W. B.; Saville, D. A.; Schowalter, W. R., *Colloidal Dispersions*; Cambridge University Press: 1989.

(6) Aoki, T. *Membranes* **2017**, *7*, 1–19.

(7) Son, S.; Takatori, S. C.; Belardi, B.; Podolski, M.; Bakalara, M. H.; Fletcher, D. A.; Fletcher, D. A.; Fletcher, D. A. *Proceedings of the National Academy of Sciences of the United States of America* **2020**, *117*, 14209–14219.

(8) Bryk, A. H.; Wiśniewski, J. R. *Journal of Proteome Research* **2017**, *16*, 2752–2761.

(9) Gautier, E. F.; Leduc, M.; Cochet, S.; Bailly, K.; Lacombe, C.; Mohandas, N.; Guillonneau, F.; El Nemer, W.; Mayeux, P. *Blood Advances* **2018**, *2*, 2646–2657.

(10) Shi, J.; Yang, T.; Kataoka, S.; Zhang, Y.; Diaz, A. J.; Cremer, P. S. *Journal of the American Chemical Society* **2007**, *129*, 5954–5961.

(11) Yang, T.; Baryshnikova, O. K.; Mao, H.; Holden, M. A.; Cremer, P. S. *Journal of the American Chemical Society* **2003**, *125*, 4779–4784.

(12) Jung, H.; Robison, A. D.; Cremer, P. S. *Journal of Structural Biology* **2009**, *168*, 90–94.

(13) Jung, H.; Yang, T.; Lasagna, M. D.; Shi, J.; Reinhart, G. D.; Cremer, P. S. *Biophysical Journal* **2008**, *94*, 3094–3103.

(14) Delaveris, C. S.; Webster, E. R.; Banik, S. M.; Boxer, S. G.; Bertozzi, C. R. *Proceedings of the National Academy of Sciences of the United States of America* **2020**, *117*, 12643–12650.

(15) Phillips, R.; Kondev, J.; Thierot, J.; Garcia, H., *Physical Biology of the Cell*, 2nd ed.; Garland Science, Taylor and Francis Group: 2012.

(16) Veatch, S. L.; Keller, S. L. *Biophysical Journal* **2003**, *85*, 3074–3083.

(17) Anderson, J. A.; Glaser, J.; Glotzer, S. C. *Computational Materials Science* **2020**, *173*, DOI: 10.1016/j.commatsci.2019.109363.

(18) Lee, H.; De Vries, A. H.; Marrink, S. J.; Pastor, R. W. *Journal of Physical Chemistry B* **2009**, *113*, 13186–13194.

(19) De Michele, C.; De Los Rios, P.; Foffi, G.; Piazza, F. *PLoS Computational Biology* **2016**, *12*, 1–17.

(20) Tan, Y. H.; Liu, M.; Nolting, B.; Go, J. G.; Gervay-Hague, J.; Liu, G.-y. *ACS Nano* **2008**, *2*, 2374–2384.
